# Surfactant-free production of biomimetic artificial cells using PDMS-based microfluidics

**DOI:** 10.1101/2020.10.23.346932

**Authors:** Naresh Yandrapalli, Julien Petit, Oliver Bäumchen, Tom Robinson

**Affiliations:** Max Planck Institute of Colloids and Interfaces, Department of Theory and Bio-Systems, Am Mühlenberg 1, 14424 Potsdam, Germany; Max Planck Institute for Dynamics and Self-Organization, Am Faßberg 17, 37077 Göttingen, Germany

**Keywords:** liposomes, lipid vesicles, surfactant-free, biomimetic, artificial cells, synthetic cells, microfluidics, bottom-up synthetic biology

## Abstract

Microfluidic-based production of cellular mimics (e.g. giant vesicles) presents a paradigm-shift in the development of artificial cells. While encapsulation rates are high and vesicles are mono-disperse compared to swelling-based techniques, current microfluidic emulsion-based methods heavily rely on the addition of additives such as surfactants, glycerol and even ethanol to produce stable vesicles. In this work, we present a microfluidic platform designed for the production of cellular mimics in the form of giant unilamellar vesicles (GUVs). Our PDMS-based device comprises a double cross-junction and a serpentine-shaped shear inducing module to produce surfactant-free and additive-free monodisperse biomimetic GUVs. Vesicles can be made with neutral and charged lipids in physiological buffers and, unlike previous works, it is possible to produce them with pure water both inside and outside. By not employing surfactants such as block co-polymers, additives like glycerol, and long-chain poly-vinyl alcohol that are known to alter the properties of lipid membranes, the vesicles are rendered truly biomimetic. The membrane functionality and stability are validated by lipid diffusion, membrane protein incorporation, and leakage assays. To demonstrate the usability of the GUVs using this method, various macromolecules such as DNA, smaller liposomes, mammalian cells and even microspheres are encapsulated within the GUVs.

## Introduction

Lipid-based vesicles have grown immense popularity in both basic research and application-oriented sciences, especially in pharmaceutics and cosmetics.^[1]^ Lipids are not only biocompatible but are also the building blocks of life-forming vesicular structures, i.e. cells. While applications of lipid vesicles in the field of health care is advancing in the form of nanometer sized liposomes,^[2]^ their usability in understanding the evolution of cells and their various biochemical and physical pathways has hit a road block due to lack of appropriate methods to form truly biomimetic cellular models. A cell mimic should fulfill the basic requirements of being lipid-based, vesicular in structure, and encapsulating the desired biomolecules such as proteins, enzymes, DNA, and even smaller vesicles as artificial organelles.^[3]^ The latter being an essential step in the emerging and accelerating field of bottom-up synthetic biology. ^[4,5]^ Currently, the most common methods include well-established electroformation and spontaneous swelling to produce lipid-based vesicular structures.^[6]^ However, the limitations of these methods prevent high and uniform encapsulation of large and charged biomolecules. Although there are reports of encapsulating biomolecules using swelling-based techniques, the reliability and reproducibility are low.^[7,8]^ In the past few years, our group and many other researchers have turned towards emulsion-based technologies to overcome these issues and precisely control the uniformity of the encapsulates.^[9–13]^ In our previous work, we showed that the inverted emulsion method can be a straightforward and reliable technique to produce basic models for cells, albeit with some drawbacks such as lack of size control, low-throughput, and its dependency on the density of the solutions used.^[9]^ Interestingly, all of the aforementioned drawbacks can be mitigated by implementing microfluidics to make lipid-based vesicles.^[14–18]^ This involves microdroplet technology to create double emulsions of water-in-oil-in-water (W/O/W) with lipids in the oil phase. Typically, the production of lipid vesicles using microfluidics (see Figure 1 (a)) involves an inner aqueous solution (IA) that is sheared by an oil phase containing lipids (LO). This results in the formation of water-in-oil (W/O) single emulsions whose interface is assembled with a monolayer of lipids, thanks to their amphiphilic nature. The single emulsion is further sheared into droplets by an outer aqueous solution (OA) to form W/O/W double emulsion. Lipids present in the oil phase assemble along both the interfaces as the oil de-wets or is exacted to form liposomes. However, a major question remains on the biomimetic properties of the lipid vesicles due to the usage of surfactants and additives in both the aqueous phases.^[14,15,19]^ For example, tri-block co-polymers made of poly(propylene)-co-poly(ethylene glycol)-co-poly(propylene) are extensively used to stabilize the liposome formation.^[16,19]^ The copolymer works by incorporating itself, more specifically, the poly(propylene) chains, into the hydrophobic region of the lipid membrane, thus altering the biophysical properties of the membrane.^[20–22]^ Furthermore, additives such as glycerol and poly(vinyl alcohol) (PVA) used to improve the viscosity of the outer and inner aqueous phases for better manipulation of fluid flow, size control and emulsion stability, also affects the membrane properties.^[14,23]^ It is very well understood that glycerol as much as 10 wt.% can completely saturate the lipid head groups in a membrane and even crosslink by acting as a bridge between phosphates present in the head groups of adjacent lipids.^[24,25]^ During the production of microfluidic liposomes as much as 15 wt.% glycerol is being employed.^[14,19]^ Furthermore, PVA polymer chains can incorporate themselves across the bilayer and in some cases enhance protein aggregation.^[26,27]^ Finally, ethanol that was used in many studies as an additive is well-known to alter the membrane properties.^[15,17,28]^ In light of these potential caveats it is important to substantially advance existing microfluidic technologies and to be able to produce lipid vesicles that can encapsulate large biomolecules in high-throughput and yet, remain biomimetic. Such biomimetic lipid vesicles can potentially be used to progress the development of the long envisioned concept of an artificial cell.^[4]^ In this work, a microfluidic device design that can be used to produce said biomimetic lipid vesicles with greater ease and in high-throughput is proposed. Furthermore, we have explored the possibilities of producing vesicles of different sizes and provided evidence of minimal to non-traceable levels of oil remaining in the membranes by tuning fluid flow rates. We demonstrate the flexibility of the method by creating membranes with neutral and charged lipids, as well formation in a variety of environments - both physiological buffers and pure water. Finally, membrane functionality and the capability of encapsulating a wide range of biomolecules within the GUVs is demonstrated.

**Figure 1.**
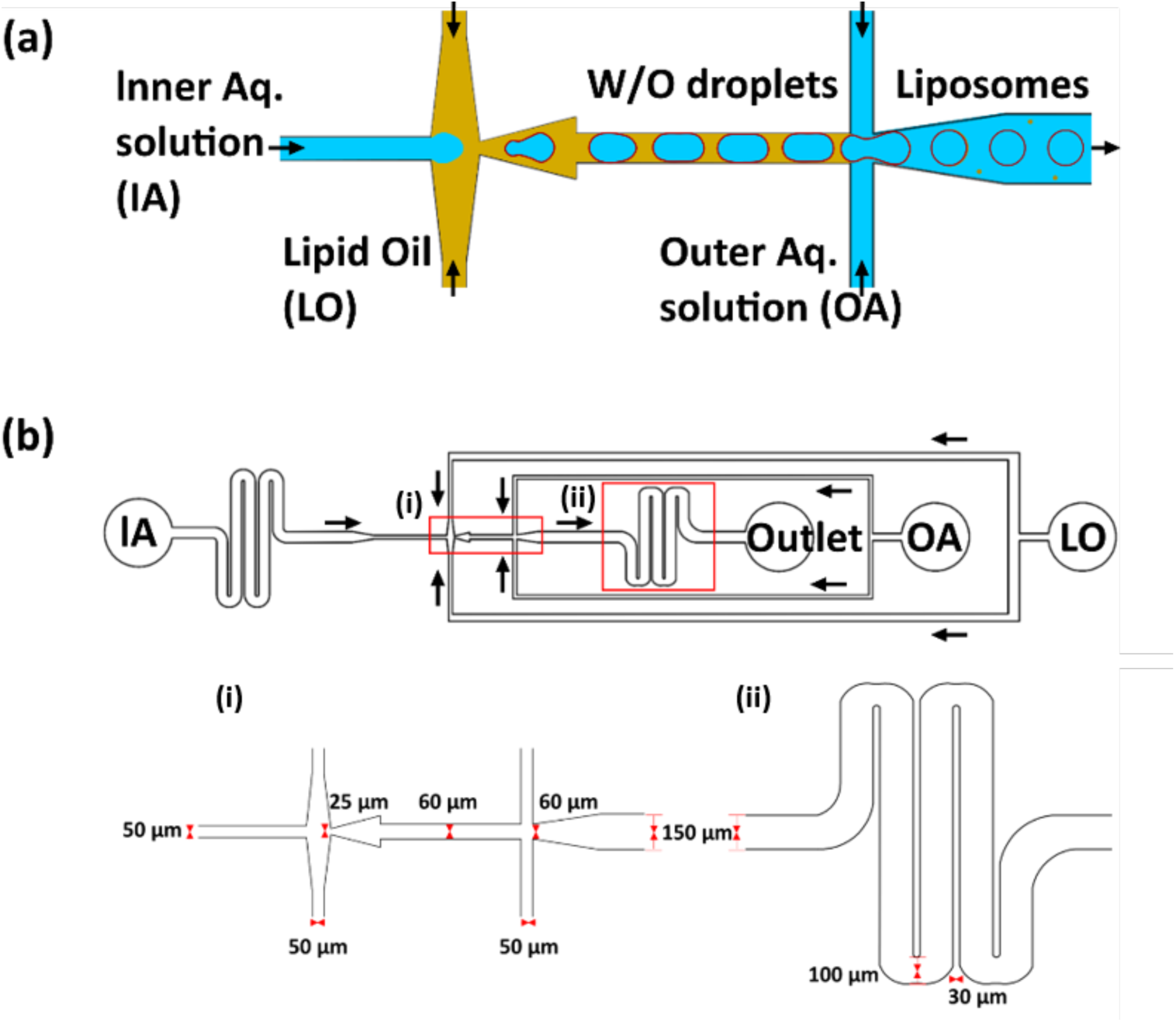
Microfluidic device design. (a) Schematic representation of liposome production using double cross-junction design and (b) design of the microfluidic chip used in this study with details on various inlets and the flow directions (black arrows) from various channels. Insets show (i) the dimensions of the two cross-junctions (ii) and the second serpentine module after the junctions.

## Results

A microfluidic design with a double cross-junction is implemented to produce liposomes in this study (Figure 1(b)). The microfluidic chip is fabricated using PDMS-based techniques (see details in Experimental section) and plasma bonded to a glass coverslip (Figure S1(a) in Supplementary information). Apart from the standard cross-junction to produce droplets, serpentine shaped buffering channels are implemented in the design to reduce the risk of fluid back flow and act as fluidic resistors. One serpentine module is implemented before the first cross-junction where the IA is sheared into droplets by the LO and a second serpentine module is implemented after the second cross-junction (Figure 1(b)). Without these modules, there is a marked decrease in flow stability and more flow discrepancies for longer periods (data not shown). Indeed, the entire production process remains stable with no requirement of flow rate adjustments over a period of approximately one hour or until the reservoirs ran out of the solution (mostly OA, due to the higher flow rate used). Apart from providing flow stability, the second serpentine module has constrictions (width is reduced from 150 to 100 µm) at every 180-degree turn, a total of four, providing increased fluidic velocities (Figure 1(b)). These constrictions impart more shear force (see Figure S1(b) for computational fluid dynamic analysis) while squeezing the double emulsions for excess oil removal (Figure S2(a)).^[14,29,30]^ The intrinsic friction provided by the serpentine module induces fast advective transport of lipids towards the interfaces to form stable membranes.^[29]^ Furthermore, the high lipid concentration in the LO substitutes the need for surfactant usage to stabilize the liposomes and also contributes to faster monolayer assembly at the interfaces. Figure 2(a) shows an example of liposomes produced by pumping MilliQ^®^ water as both the IA and OA and 1-palmitoyl-2-oleoyl-sn-glycero-3-phosphocholine (POPC) (5mg/mL) in 1-octanol as LO. To achieve this, initially the surface functionalized chip (see details in Experimental section) is wetted by pumping OA through the channels, followed by LO and IA. This strategy prevents the oil from eroding the surface coating of the OA-Outlet channel (a layer-by-layer self-assembled polyelectrolyte). Upon introduction of all the three fluids into the chip, flow rates (via pressure) are adjusted to produce liposomes of a specific diameter. In this case, fluid flow of LO is set to 44 mbar while IA to 50 mbar and OA to 57 mbar of pressure (see SI for Video 1). The resultant double emulsion formation rate is typically 3.2 ± 0.6 kHz. The high flow rates implemented at the second cross-junction are necessary to shear the W/O droplets and form very thin shells of the oil-phase (W/O/W). The thinness of the shell equates to lower amounts of 1-octanol and thus, faster removal with the help of the shear modules. The leftover oil residue (if any) easily dewets from the double emulsion (Figure S2(b)). The final lipid vesicles, produced with only water both inside and outside, and with a diameter of 120 ± 5 µm are shown in the Figure 2(b). To the best of our knowledge, these are first giant lipid vesicles produced with pure water both inside and outside exhibiting very high size homogeneity and yield. It is important to specify that no surfactants or additives are used in the production process in order to retain the biomimetic aspects of the resulting membranes. The lack of thickening agents like PVA or glycerol, requires the design to contain narrow channels and fluidic resistors to allow for easy tuning of the vesicle sizes. The device allows for the production of vesicles not only with a narrow size distribution but with tunable diameters (at least an order of magnitude range) unlike previous studies on microfluidic vesicle production.^[14,23]^ Figure 2(c) depicts the histograms of vesicles with diameters i.e., 13 ± 2.1 µm with a relative standard deviation (RSD) of 15.8%, 60 ± 1.9 with RSD of 3.2 %, 87 ± 1.6 with RSD of 1.8%, 101 ± 2.7 with RSD of 2.7% and 130 ± 4.2 µm with RSD 3.2%. Note that the RSD value of the smallest set of liposomes produced is approximately three times higher than the vesicles of larger diameter in size, which is most likely due to the very high flow rates that have to be employed to yield smaller diameters and therefore reduced control at the second cross-junction/vesicle forming junction. If need be, this can be avoided by employing high molecular weight polyethylene glycol (PEG) in the outer aqueous solution (data not shown). Furthermore, we have tested the possibility of producing negatively charged lipid vesicles using this microfluidic device. Negatively charged membranes provide an opportunity to study the interaction and assembly of many positively charged membrane proteins, as well serving as bacterial membrane models. Figure 2(d) shows confocal images of GUVs containing 1,2-dioleoyl-sn-glycero-3-phospho-L-serine (DOPS) in their membranes. The homogeneity in the size of the charged vesicles produced (67 ± 2 µm with RSD of 2%) and the encapsulation uniformity (here calcein dye) suggests the versatility of the device compared to standard technique like electroformation where it is not possible to efficiently produce charged lipid vesicles.

**Figure 2.**
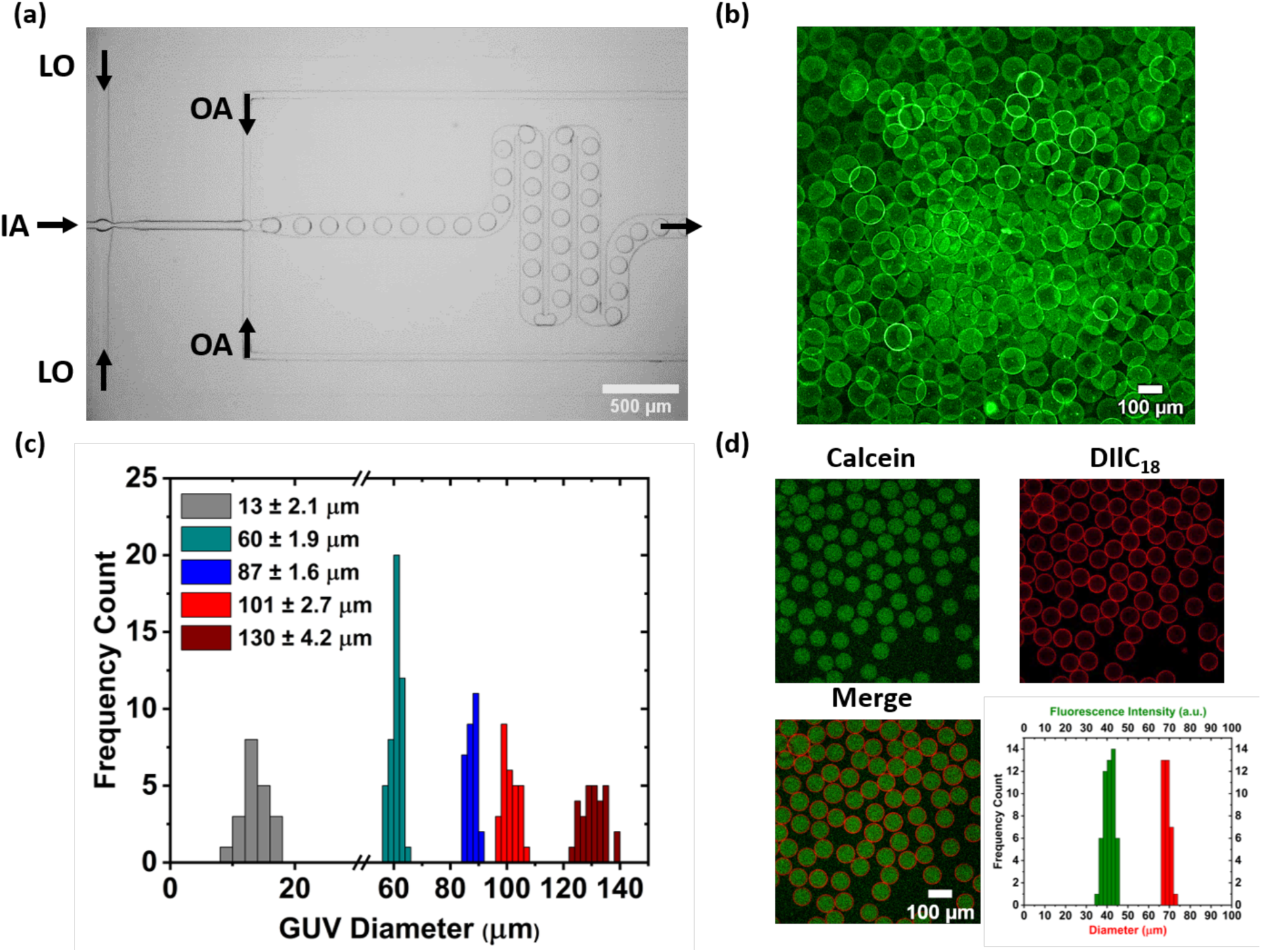
Production of monodisperse additive-free pure-lipid liposomes. (a) The production process using the microfluidic double emulsion device with 50 mbar at the IA (inner aqueous solution), 57 mbar at the OA (outer aqueous solution) and 44 mbar at the LO (lipid oil). (b) Confocal fluorescence image of the POPC GUVs obtained with only water as the IA and OA. (c) Size distributions of the liposomes for different flow rate configurations, showing high size monodispersity over a wide range. (d) Confocal fluorescence images of negatively charged GUVs with lipid composition POPC (79.5 mol%), DOPS (20 mol%), DIlC18 (0.5 mol%) containing calcein (10 µM) together with the encapsulation efficiency (100%) and uniformity (RSD 6.2%) as well as their size homogeneity (67 ± 2 µm, RSD 2%).

To keep the vesicles biomimetic, it is not only important to use surfactant-free and additive-free solutions that can adversely affect the membrane biophysical properties, it is also important to quantify and validate the membrane properties systematically. In this respect, Figure 3 shows the gradual decrease in the oil thickness present in the immediately formed double emulsion to finally achieve optically non-traceable levels. Figure 3a depicts wide-field images of the double-emulsions produced using the microfluidic device described earlier. The flow rates of the OA is gradually increased while the flow rates of the IA and LO are kept constant. By designing the OA channel width (50 µm) to be one third the width of outlet (150 µm), it is possible to shear W/O droplets with exceptional control to form W/O/W emulsions. Gradual increase in the flow rate of the OA channel made a difference of oil thickness from 35 ± 2 µm in diameter to negligible levels (Figure 3(b)) (see SI for Video 2). To further ascertain the biomimetic nature of the liposomes, fluorescence recovery after photo bleaching (FRAP) was performed. Lipid lateral diffusion is a signature characteristic that defines the purity of the membrane and can reveal impurities if the measured diffusion times differ from control values.^[31–33]^ The diffusion times are determined by photobleaching the labeled lipids and observing the recovery of fluorescence in the bleached region over time. Using the information, it is possible to calculate the mobile fraction and hence the diffusion coefficient of the bleached lipid (see Experimental section for details on the method). FRAP experiments are performed on liposomes containing 300 mOsm sucrose that are produced using both microfluidics and electroformation (see Figure 4). The latter being the control as oils are not used. In both cases, the lipid composition is set as 99.5% POPC and 0.5% NBD PC. Figure 4(a) and 4(b) shows the average fluorescence intensity recovery curves of the liposomes after photobleaching. Typical images of the bleached region of the liposomes along with snapshots before and after the exposure are also provided. From the fitting of curves, the diffusion coefficients of NBD PC in electroformed and microfluidic liposomes were determined to be 7.5 ± 1.8 and 8.5 ± 4 µm^2^/sec, respectively. These values suggest that the lipid lateral diffusion in membranes is similar for both electroformed and our microfluidic liposomes. Furthermore, the diffusion coefficient observed in these measurements are in agreement with previous literature values.^[31,34]^

**Figure 3.**
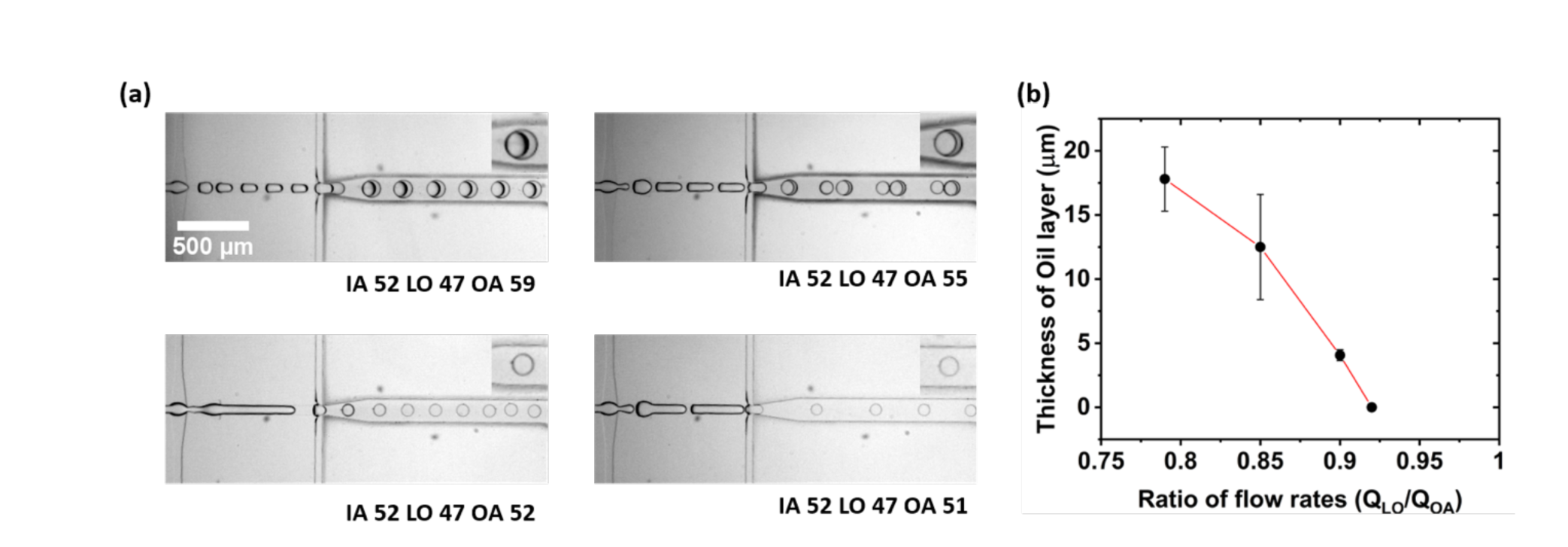
Manipulation of flow rates for reduced oil content and ultra-thin shells. (a) High-speed camera images of the microfluidic liposomal production process to reduce the oil thickness. Inserts: enlarged images of individual double emulsions. From top left to bottom right corner, the oil thickness has been reduced by gradually tuning the flow rates of the OA (pressures for each channel are given in mbar). (b) Plot showing the thickness of the oil layer that is reduced from 35 ± 2 µm to 24 ± 4 to 8 ± 0.5 to finally optically non-detectable amounts.

**Figure 4.**
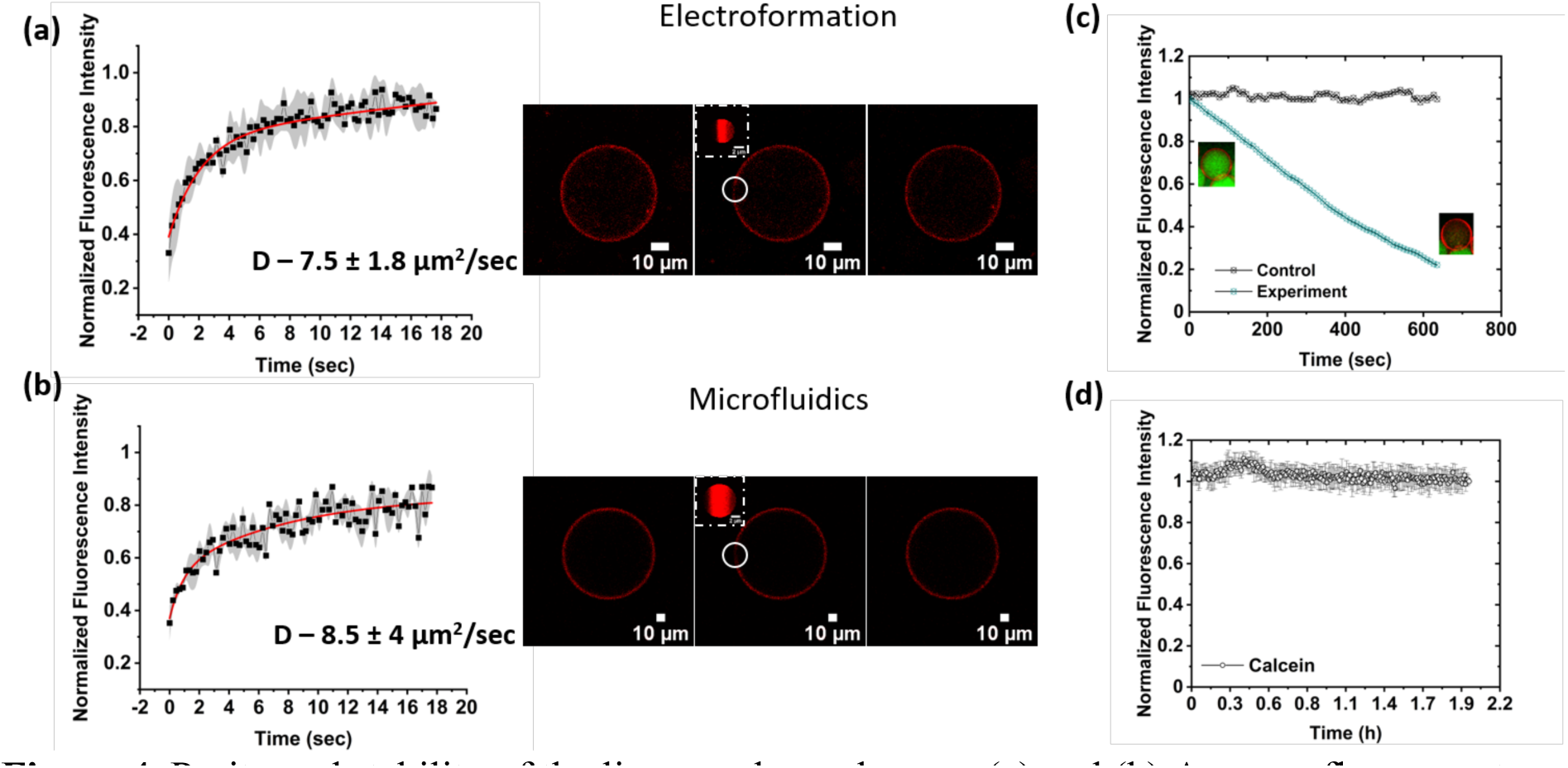
Purity and stability of the liposomal membranes. (a) and (b) Average fluorescent recovery curves of at least five liposomes produced using electroformation and microfluidics along with the confocal cross-sections of before and after photobleaching. (c) Plot depicting the calcein dye-leakage assay using α-hemolysin protein insertion with a gradual decrease in mean luminal fluorescence intensity over time (n > 10). (d) Plot showing the vesicle stability over-time with no decrease in the mean calcein fluorescence intensity over time (n > 10).

To further assess the quality of the liposomal lipid membrane produced in this work, a membrane protein induced leakage assay was performed. Using α-hemolysin, a membrane spanning heptamer, calcein dye leakage from the lumen of the liposomes into the outer solution was monitored. Microfluidic liposomes were produced with 20 µM calcein in the IA, placed on a bovine serum albumin (BSA) coated coverslip, and flushed with α-hemolysin solution (final concentration of 2.5 µg/mL). The leakage profile of the calcein dye is plotted in Figure 4(c). Within a span of 10 min, calcein leakage from the lumen of the liposomes is observed while the fluorescence intensity of the control (no membrane pores) remained intact during the entire period. Not only does this finding demonstrate the functionality of the membranes but also their unilamellarity. In addition to this, the overall stability of the calcein filled GUVs were analyzed for a period of 2h. Figure 4(d) shows that the calcein intensity inside the lumen of the liposomes remained constant and the vesicles are stable when there is no external influence. Collectively all the studies performed: optically undetectable oil traces (Figure 3), FRAP experiments to confirm the unhindered lipid lateral diffusion (Figure 4(a) and 4(b)), membrane protein induced dye leakage assay (Figure 4(c)), vesicle stability studies (Figure 4(d)), and all without the use of surfactants or additives, supports the biomimetic pure-lipid nature of the liposomes produced with our microfluidic technique.

We have tested the applicability of the microfluidic device and also ascertained its robustness to produce artificial cells/carriers of various kinds. The possibility to incorporate large biomolecules is the major promise of using emulsion-based methods and microfluidics for their high-throughput capabilities. In Figure 5, experiments were conducted to proof this by incorporating a range of large molecules from plasmid DNA to microspheres. Firstly, as a proof-of-concept, IA containing circular plasmid DNA and EvaGreen^®^ dye was used to make liposomes with water as the OA (Figure 5(a) and SI for Video 3). The produced liposomes are not only homogeneous in size, but also have uniform green fluorescence intensity across all of the analyzed luminal cross-section and not on the lipid membrane (see SI for Figure S3). The uniformity shown here proves that the lipid membrane and the lumen of the vesicle are free of oil residues. This result is a step towards building an artificial cell – by adding ingredients required for translation and transcription like in prokaryotic cells. As a follow-up to the concept of constructing artificial cells, we have incorporated HEPES buffer containing small unilamellar vesicles (SUVs) inside these giant liposomes to mimic eukaryotic cell architecture (Figure 5(b)). SUVs are highly robust small vesicles in the range of 50 nm in diameter. Protocols for producing and encapsulating desired materials inside these SUVs and even LUVs (large unilamellar vesicles) are very well established.^[2]^ We present the microfluidic device and the associated result (Figure 5(b)) as a potential way to study the evolution of prokaryotic cells to eukaryotic cells, more specifically compartmentalization and their role in organized decentralization within cells. Note that the high-throughput nature of the microfluidics is evident from the large populations of liposomes that can be seen in the Figure 5(b) and robustness of the technique in using HEPES or other buffered solutions with ease, unlike standard techniques like electroformation. A step further to this would be to encapsulate cells within the liposomes (Figure 5(c)). Lipid-based vesicles present a natural environment for the cellular growth, considering that cells interact with other cells in tissues or biofilms. Recently, biofilms have been made within droplets that can be used for understanding biofilm growth, expansion, and even high-throughput screening.^[35,36]^ Lipid vesicles that can be produced with ease using PDMS-based microfluidics will be a boon to either single cell growth studies or even development of organoids and biofilms for drug discovery. PDMS-based devices provide an additional advantage as the produced vesicles can be subsequent captured and analyzed in the same device.^[37]^ The data in Figure 5c demonstrates the ability to encapsulate fibroblast cells using this device. One other interesting aspect of utilizing microfluidics to produce lipid vesicles is the possibility to incorporate very large molecules, even non-biological foreign bodies. In Figure 5(d), we incorporated large styrene microspheres of ~20 µm diameter inside the microfluidic liposomes. While it was not possible to incorporate microspheres inside every liposome, due to not being dispersible in aqueous solutions unlike above-mentioned examples (plasmid, liposomes and cells), approximately 30% contained microspheres. Considering the high-throughput nature of the production process, 30% will still result in high number to study the interaction of non-degradable, environmentally toxic materials and their digestion.

**Figure 5.**
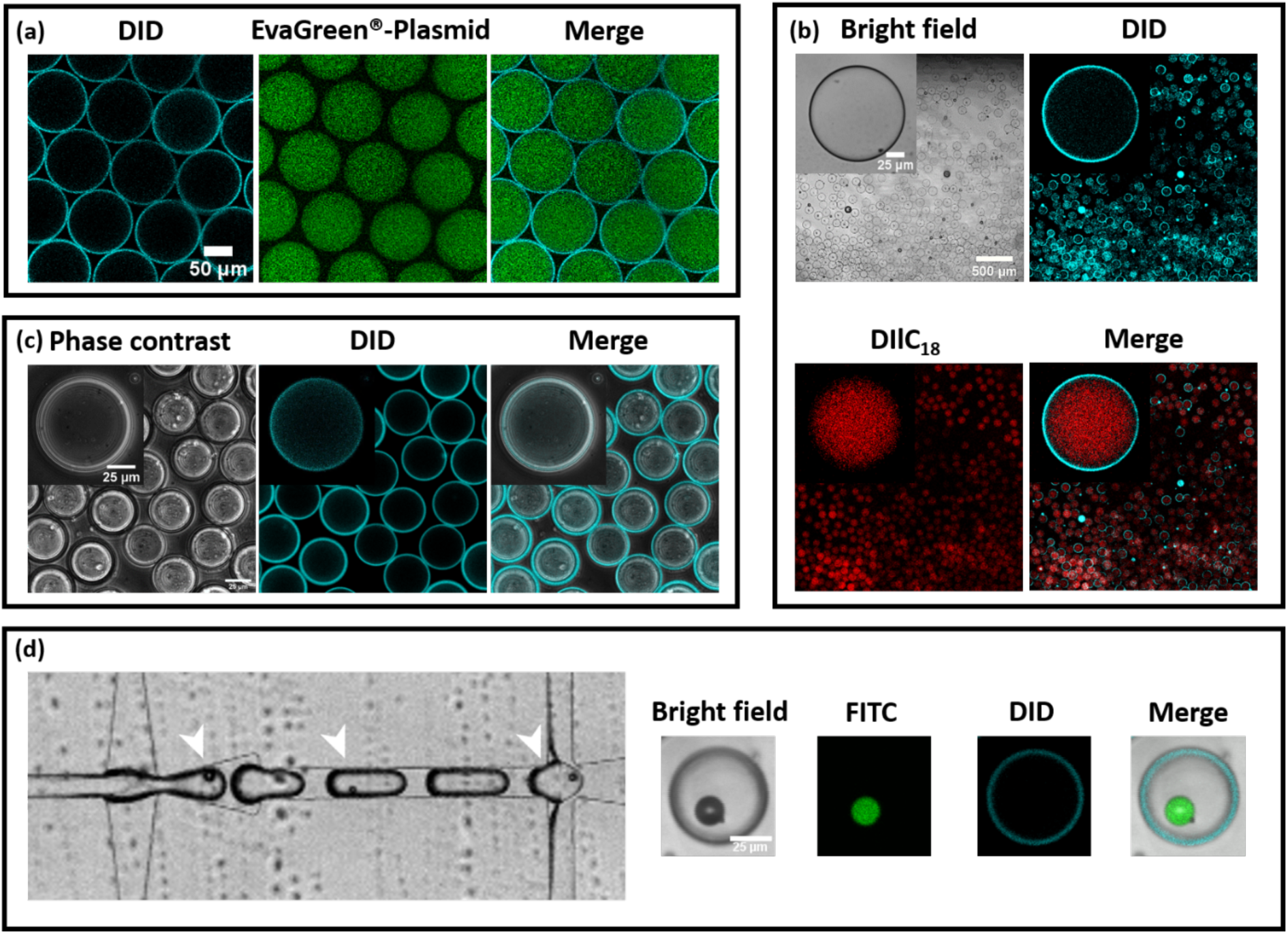
Microfluidic production of artificial cells (a) encapsulating EvaGreen^®^-plasmid DNA, (b) showing compartmentalization with SUVs, (c) encapsulating fibroblast cells along with the culture media, and (d) encapsulating microplastics such as styrene microspheres.

## Conclusions

Taken together, these results show that the microfluidic device proposed in this work can be used for making double emulsions in general, and GUVs in particular, with great ease and in a reliable manner. Since the device itself is made from standard PDMS-based methodologies, reproduction and usage are facile as well offering great versatlity. Monodisperse size control of the GUVs is possible within a wide range from ~20 µm to 120 µm in diameter. The membranes can be composed of neutral as well as charged lipids (binary mixtures), in a variety of solutions (pure water, physiological buffers, culture media), and in the absence of the surfactant or additives. The resulting biomimetic properties of the liposomes and their stability have been validated using both optical and fluorescence based techniques such as FRAP and dye leakage assays. Applicability of the device in producing artificial cells is thoroughly explored by encapsulating small molecules to large living and non-living components such as cells and microspheres. The promise of emulsion-based technologies is immense and microfluidic device design and methodologies employed here will help progress the field of artificial cells considerably.

## Supporting Information

Supporting Information is available.

## Acknowledgements

This work is part of the MaxSynBio consortium which is jointly funded by the Federal Ministry of Education and Research of Germany and the Max Planck Society. The authors would like to thank David Gonzales and Dora Tang for gifting the DNA plasmid, Christine Pilz-Allen for supplying the fibroblast cells, and Reinhard Lipowsky for financial support.

---

Artificial cells are key to understanding the sophisticated functions of biological cells. Microfluidics paves the way to produce them as micron-sized lipid vesicles in a high-throughput and tunable manner. The device presented in this work is able to create truly biomimetic lipid membranes free from additives or surfactants, taking us one-step closer towards fully functioning artificial cells.

**Keyword** Artificial cells

Naresh Yandrapalli, Julien Petit, Oliver Bäumchen, Tom Robinson*

Surfactant-free production of biomimetic artificial cells using PDMS-based microfluidics

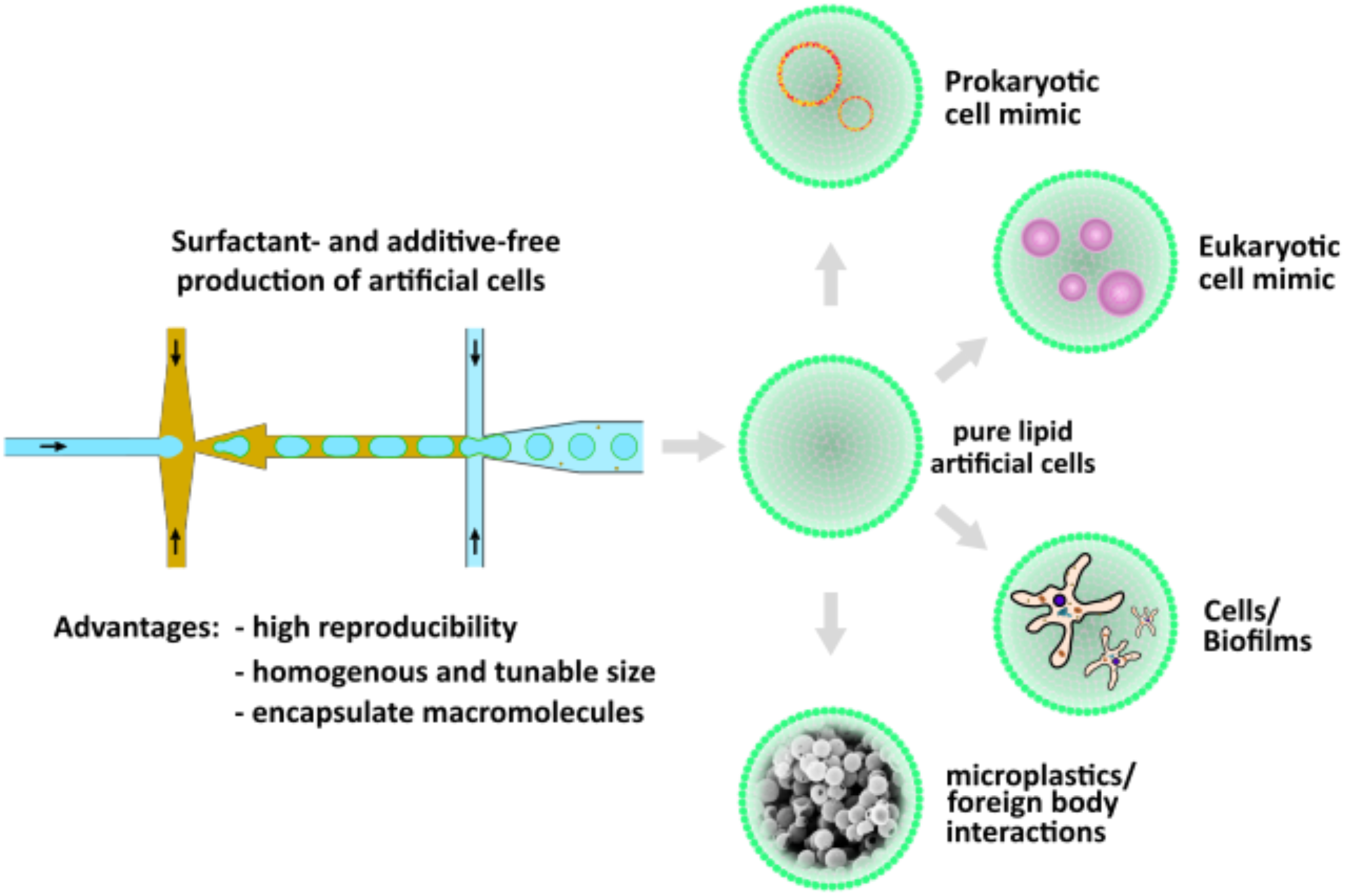

## Notes

### Competing Interest Statement

The authors have declared no competing interest.

