## Supplemental for "Surfactant-free production of biomimetic artificial cells using PDMS-based microfluidics"

### Materials

All materials were used as purchased unless noted otherwise. 1-octanol (99 %, Sigma Aldrich), (97%, Sigma Aldrich). 1-Palmitoyl-2-oleoyl-sn-glycero-3-phosphocholine (POPC), 1,2-dioleoyl-sn-glycero-3-phospho-L-serine (DOPS) and 1-palmitoyl-2-{6-[(7-nitro-2-1,3-benzoxadiazol-4-yl)amino]hexanoyl}-sn-glycero-3-phosphocholine (NBD PC). was obtained from Avanti polar lipids. 1,1′-Dioctadecyl-3,3,3′,3′-tetramethylindotricarbocyanine perchlorate ($DiIC_{18}$ (3)) and DID ($DIlC_{18}$ (5)) and calcein were purchased from ThermoFisher Scientific. Polydimethylsiloxane (PDMS) and curing agent were obtained as SYLGARD®184 silicone elastomer kit from Dow Corning. 1H,1H,2H,2H-Perfluorodecyltrichlorosilane was purchased from abcr GmbH. Poly(diallyldimethylammonium) chloride (PDADMAC) and poly(sodium 4-styrenesulfonate) (PSS) were obtained from Sigma Aldrich. SU8 2050 (Microchem Inc.), Silicon wafer (Siegert Wafers), SU8 developer solution (Microchem Inc.). Indium tin oxide (ITO) glass slides are from Präzisions Glas & Optik GmbH. Glucose, sucrose and chloroform were obtained from Merck. HEPES and PBS were purchased from Sigma-Aldrich. Dulbecco's Modified Eagle Medium (DMEM) was purchased from ThermoFisher Scientific. Mini Extruder and polycarbonate membranes were purchased from Avanti polar lipids, Inc. Fluoresbrite® YG Microspheres purchased from Polysciences, Inc. Microfluidic lipid vesicle production was achieved using a MFCS™-EX pressure pump with associated 2 mL reservoirs for various solutions from Fluigent, Inc.

### Experimental

#### Microfluidic device fabrication

The PMDS-based microfluidic device fabrication was performed using soft-photolithography as described previously.[1] Master moulds were prepared on 4″ silicon wafers using spin coating (model no. WS-650MZ-23NPPB, Laurell Tech. Corp.) SU8 2025 (Microchem Inc.) to a height of 80 µm. Following the coating process, a pre-baking step was performed before UV-light exposure through a film mask with the requisite design, onto the SU8 coated silicon wafer for a duration of

8 sec (MicroLithography Services). A post baking step was performed before SU8 development process. SU8 development was performed by gently washing the wafer in developer solution (Microchem Inc.) for 3 minutes. Finally, the Si-wafer was hard baked for a period of 30 min at 200 ºC. Then the prepared master moulds were silanised overnight (50µl of 1H,1H,2H,2H-perfluorodecyltrichlorosilane) in a desiccator. PDMS-based microfluidic chips were produced by heat curing (90 ºC for three hours) PDMS with curing agent mixture (10:1) on the master mould Si-wafer. Cured PDMS was peeled and cut into individual chips. Holes were punched at respective inlet and outlets using a 1 mm biopsy puncher (Kai Europe GmbH) before bonding the PDMS to glass coverslips. At the end, bonding was performed using an air plasma treatment (Plasma Cleaner PDC-002-CE, Harrick Plasma) at 600 mbar for 1 min. The microfluidic devices were heated for 2 h at 60°C to help with the bonding process and render the PDMS back to hydrophobic after the plasma treatment. A photograph of a final assembled device can be seen in the Figure S1a.

W/O/W emulsion production requires the surface of the outer channel (from OA inlet to outlet, see Figure 1b in the main text) to be hydrophilic. Unless the outer channel is wettable by aqueous medium, W/O/W type of double emulsion will not form. This is because the material used in this study to make the microfluidic chip, PDMS, is hydrophobic in nature and is not wettable by aqueous medium. For the outer aqueous solution (OA) to be able to wet the PDMS surface, the channel was treated with series of chemical reagents. Channels carrying OA towards the outlet (see Figure 1b) was subjected to an initial cleaning step by flushing $HCl:H_2O_2$ (1:2) for 30 sec. This renders the chip surface negatively charged for polymeric solution of positively charged polymer (2 wt.% PDADMAC) to form layers after flushing the solution for 2 min. Following this, a negatively charged polymeric solution (5 wt.% PSS) was flushed for the same duration to yield hydrophilic OA outlet channel. After every step, mention above, MilliQ® water was flushed in for a minimum of 30 sec to remove excess chemical reagents. Thus coated channels remained hydrophilic for more than a week. Importantly, the entire coating process takes up only 6 minutes to complete and the device is ready to use thereafter.

**Liposome production: microfluidics, electroformation, and extrusion**

A double cross-junction chip design was used to produce the double emulsions (to eventually form GUVs). To make W/O/W double emulsion, the inner aqueous solution (IA) containing MilliQ®, EvaGreen®-plasmid DNA (equimolar mixture), calcein, SUVs, cells or styrene microspheres was passed through the first cross-junction to be sheared into aqueous droplets by 1-octanol with 5mg/mL total lipid concentration of 99.5 mol% POPC and 0.5 mol% DID/$DIlC_{18}$/NBD PC). Following this, the oil phase (LO) carrying aqueous droplets was further sheared to produce double emulsion at the second cross-junction by the outer aqueous (OA) solution of MilliQ® water/300 mOsm Glucose solution/HEPES buffer (20 mM). Electroformed GUVs were produced using indium tin oxide (ITO) coated glass plates. 15 µL of 2mg/mL total lipid concentration of 99.5 mol% POPC and 0.5 mol% NBD PC was smeared over the ITO surface side of the glass plates and dried using a nitrogen gun and further in a desiccator at low pressure conditions for 45 minutes. A chamber was created with both the lipid-coated surfaces facing inwards using a Teflon spacer. This chamber was filled with 300 mOsm sucrose solution and sealed. Electric input was delivered to the ITO sides of the glass plates through copper tapes connected to AC-input generator at 10 Hz and 2 Vp-p for 2 h. After the production, glass plates were finger-tapped for the release of the GUVs into the sucrose solution before they are collected in an Eppendorf® tube. The 50 nm size small unilamellar vesicles (SUVs) were produced using hydration and extrusion method. 24 mM total lipid concentration of 99.5 mol% POPC and 0.5 mol% DIIC18 in chloroform were dried on to a glass vial using nitrogen gun and in desiccator at low pressure conditions for 45 min. This was followed by hydration of the lipid layer with 1 mL of warm HEPES buffer (37 °C). Hydration was allowed to take place overnight at room temperature. The produced multilamellar vesicles were subjected to freeze-thaw cycle using liquid nitrogen and a hot water bath. Further to this, multilamellar vesicles (MLVs) were made into SUVs using a lipid extruder fitted with 50 nm polymeric filter. After 21 extrusion cycles, the SUVs produced were used in microfluidic production to make compartmentalized liposomes.

**Cell culture**

Fibroblast cells were cultured in DMEM media at 37 °C and retrieved using trypsin enzyme from petri dishes. The suspension is directly used as IA to encapsulate cells and to therefore demonstrate the possibility to encapsulate cells within liposomes produced using microfluidics.

**Fluorescence Recovery After Photobleaching (FRAP)**

The fluorescence image sequences of the liposomes were acquired using the 488 nm line of an Ar+ laser at a very low power to avoid photobleaching. After 2.5 s, regions of interests (ROI), of 6 µm radius were rapidly photobleached ($t < 60$ ms) at maximal laser power. Fluorescence recovery was monitored for ~ 20 s. The recovery curves were obtained as explained in a previous work.[2] In order to correctly estimate $F_0$ (the fluorescence intensity immediately after the end of the bleach) and $F_{18s}$ (after 18 s), the curves were fitted as in ref. 2. The normalized fractional recovery (NFR) is defined as:

$$NFR = \frac{F_{18s} - F_0}{1 - F_0}$$

Following this, a non-linear exponential curve fitting was performed (OriginPro, OriginLab Corp.) on the fractional recovery curve from each data set to calculate the halftime ($t_{half}$) of recovery. Using the $t_{half}$ obtained, the diffusion coefficient for the lipid DOPC-NBD was determined using:

$$D = {0.25\omega^2}/{t_{half}}$$

where D is the diffusion coefficient and ω is the radius of the bleach spot.

**Dye-leakage assay**

The leakage of calcein dye from the lipid bilayer has been used in earlier works as a tool to investigate the functionality and unilamellarity of model lipid membranes.[1,3] We have tested the liposomes produced in this work for leakage both in the presence and absence of α-hemolysin as a pore-forming protein. Liposomal vesicles were produced with 20 µM calcein solution as the IA. To which, 2.5 µg/mL final concentration of α-hemolysin was added. The graph shown in the Figure 4c presents the recording obtained after 10 minutes of incubation.

### Microscopy

The produced liposomes which were collected in Eppendorf® tubes were pipetted onto BSA coated (2 mg/mL for 30 min at 37 °C) glass coverslips and sealed using SecureSeal™ image spacers (Sigma Aldrich). A MicroLab 310 (Vision Research Inc.) high-speed camera fitted to an Olympus IX73 microscope was used to acquire images of the liposome production at full frame and with ~3000 frame rate. Liposomes produced were also visualized using confocal microscopy (Leica TCS SP8, Leica Microsystems Inc.). For image acquisition, ex488/em499-540 nm for EvaGreen®, NBD PC and Calcein, ex551/em565-610 nm for $DIlC_{18}$, and ex633/em645-680 nm for DID wavelengths were used. Data produced were treated and analyzed using Image J software.

### Computational Fluid Dynamics

To prove that constricted curves can result in increased forces, a computational fluid dynamics (CFD) simulation was performed using FEATool Multiphysics for MATLAB®. Using a 2D replication of the design used in this work, the first constriction of the serpentine channel region was drawn with the built-in geometry tools. Following this, a 2D grid was generated and refined using Gmsh mesh builder. The Navier-Stokes Equation (for incompressible fluids) was used to simulate the fluid flow.

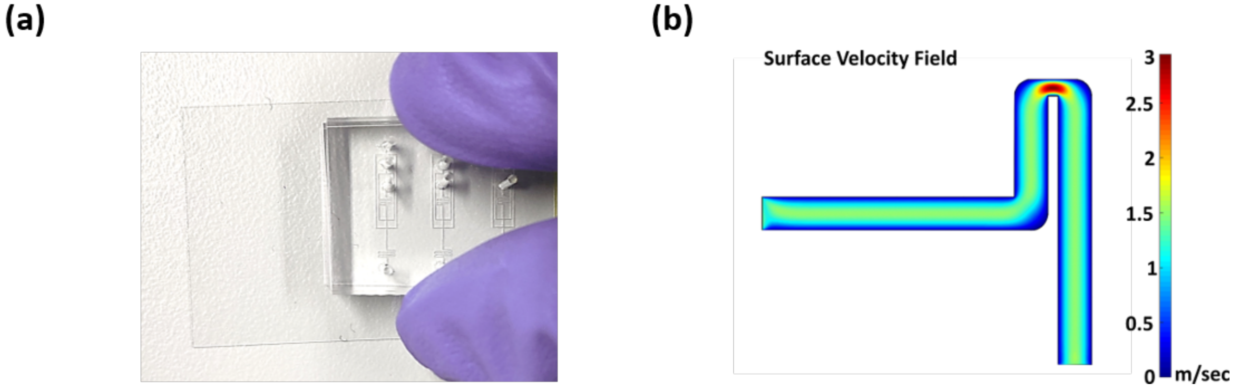


Figure S1. (a) Photograph showing the final fabricated microfluidic chip bonded to a glass coverslip. Each chip has six copies of the design for repeat experiments and convenience. (b) CFD simulation in 2D of the first constriction of the serpentine module present in our microfluidic design.

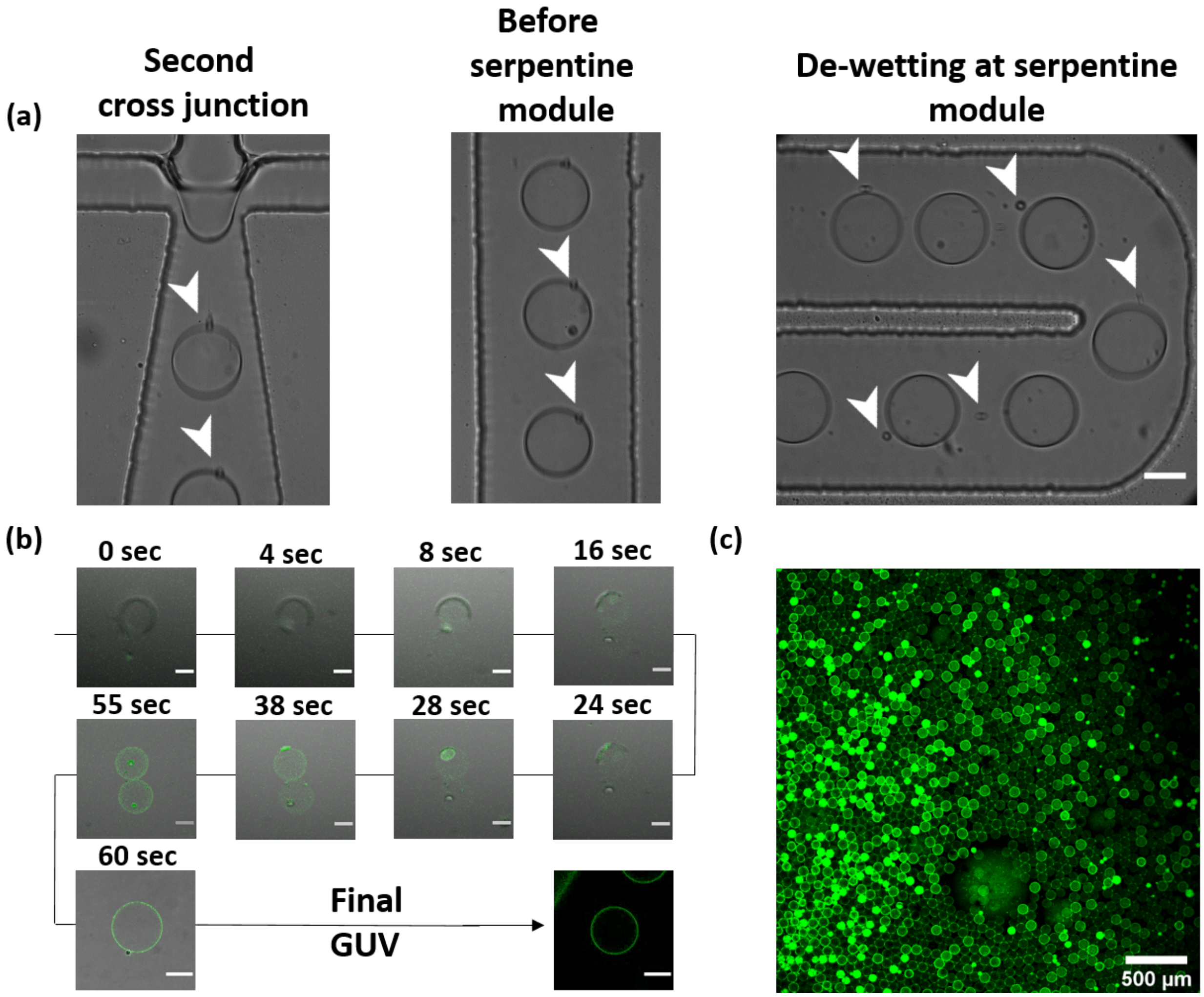


Figure S2. Dewetting process to form microfluidic lipid vesicles. (a) Snapshots of vesicles forming in the microfluidic channels, starting with the formation of double emulsion at the second junction and spontaneous 1-octanol dewetting and pinch-off - enhanced by the serpentine module (arrows showing the oil droplets). Scale bar corresponds to 50 µm). (b) Snapshots of the left-over oil residues, in some cases, dewetting and pinching-off from individual vesicles in the solution. Scale bar corresponds to 50 µm. (c) Wide field of view (x5 objective) of large number of vesicles produced using the microfluidic device along with the dewetted oil droplets (bright green spots).

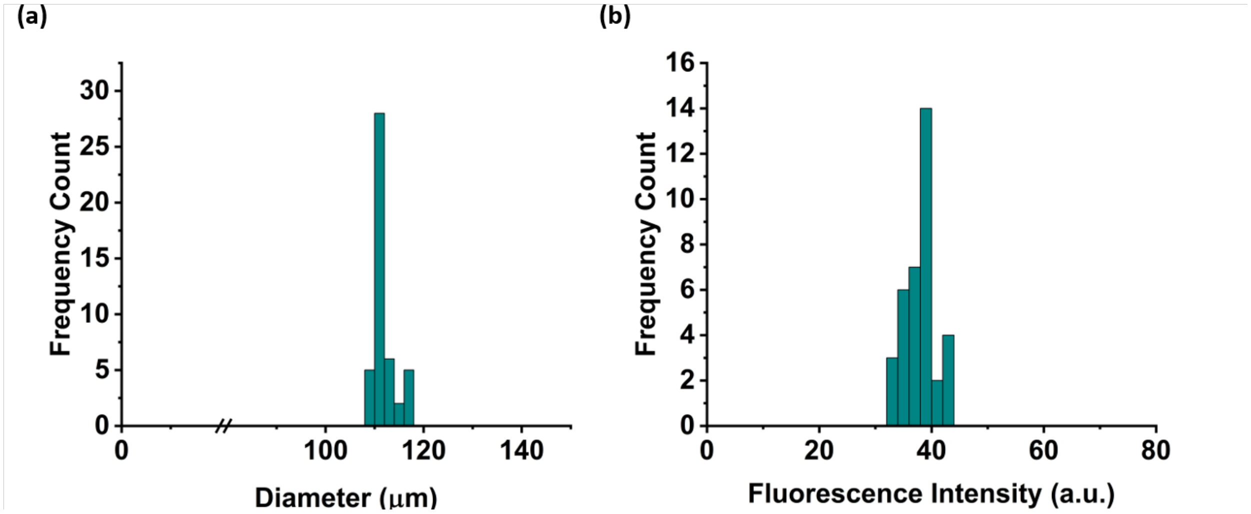


Figure S3. Homogeneous size distribution of the liposomes produced by our microfluidic technique and its encapsulation efficiency. (a) The monodisperse nature of the microfluidic GUVs containing EvaGreen®-plasmid DNA, 110 ± 4 μm with RSD of 2.5% and (b) the EvaGreen®-plasmid DNA fluorescence intensity among various GUVs produced using this technique, showing a very narrow distribution.

**Legends for Supporting Videos**

Video 1

Microfluidic production of liposomes with MilliQ® water in IA as well as in OA and POPC lipids in LO at 25 fps (IA-50 mbar, LO-44 mbar, and OA-57 mbar).

Video 2

Video showing microfluidic production of liposomes. On the right-side liposomes with thick oil layer (IA-52 mbar, LO-47 mbar, and OA-59 mbar) and the left-side with optically invisible oil layer thickness (IA-52 mbar, LO-47 mbar, and OA-51 mbar) produced by carefully altering the flow rates.

Video 3

Microfluidic production of liposomes with EvaGreen®-plasmid DNA mix as IA (51 mbar), MilliQ® water as OA (100 mbar) and POPC lipids in LO (47 mbar).

Video 4

Time-lapse video of styrene microsphere interaction with lipid membrane. At the end of the video it is evident that the microsphere has affinity towards the lipid membrane and resulted in binding.
